# Pharmaceuticals residues lead to enrichment of Human, Animal, and Plant Pathogens in Wastewater

**DOI:** 10.1101/2025.11.05.686723

**Authors:** Marcel Suleiman, Gaofei Jiang, Alexandre Jousset

## Abstract

Wastewater forms a reservoir for diverse microbial pathogens, posing significant risks to public health and aquatic ecosystems. They further contain several pharmaceutical residues that can induce profound shifts in the wastewater microbiome. In this study, we assessed whether common non-antimicrobial pharmaceuticals such as pain killers and blood pressure regulators impact wastewater pathogen profiles. We spiked wastewater with caffeine (stimulant), atenolol (beta-blocker), paracetamol (analgesic), ibuprofen (anti-inflammatory) and enalapril (ACE-inhibitor), and monitored microbiome 16S rRNA gene profiles. We then matched species composition to a comprehensive database of human, animal, and plant pathogens. All pharmaceuticals significatively increased abundance and biodiversity for zoonotic, plant, and animal pathogens. These findings highlight that pharmaceutical contamination poses a biohazard risk by fostering pathogen growth. We call for a paired, continuous monitoring of chemical and biological pollutants and a more stringent removal of pharmaceutical residues from wastewater.

## Introduction

While organic pollutants in wastewater have recently been recognized as bioactive compounds with potential direct impacts on human health, the role of these pollutants in influencing the growth and distribution of pathogens in wastewater treatment plants (WWTPs) is still poorly understood. Microbial pathogens in wastewater represent a significant public health and environmental concern [1]. Wastewater harbors hundreds of harmful bacterial species capable of causing diseases in humans, while also threatening livestock, wildlife, and agricultural production[2]. Polluted wastewater therefore functions as a reservoir for a wide range of pathogens, encompassing human, zoonotic, and plant-associated microorganisms [3]. Understanding how wastewater pollution shapes pathogen prevalence and diversity is critical for developing effective monitoring and management strategies to safeguard both public health and aquatic ecosystems. In this study, we use pharmaceuticals as model pollutants. Pharmaceutical residues are among the most prevalent pollutants in wastewater worldwide, with many molecules being recalcitrant and only partially removed during wastewater treatment. They persist in effluents discharged into the environment, with potential cascading effects on ecosystem health [4, 5]. Importantly, even pharmaceuticals without classical antimicrobial activity—such as the painkiller paracetamol, the beta-blocker atenolol, and the anti-inflammatory ibuprofen—can have a strong impact on microbial communities [6]. In a previous work, we demonstrated that such compounds can markedly shift the composition and dynamics of wastewater microbiomes [7]

The present study investigated whether pharmaceutical pollution could drive an increase in biohazard risk by promoting pathogen abundance and diversity in wastewater. We reanalyzed our previously generated dataset [7] using the MBPD pipeline [8] to quantify the prevalence and biodiversity of human, zoonotic, and plant pathogenic bacteria in relation to specific pharmaceutical exposures. We expected that pharmaceutical pollutants would impact pathogen density, that these changes would be driven by the selective growth of specific bacterial taxa, and that the magnitude and nature of the effects would vary depending on the type of pathogen. By exploring the interplay between pollutant identity and pathogen distribution, this study provides new insight into the complex mechanisms through which bioactive pollutants shape microbial risk in aquatic environments.

## Material and methods

### Dataset

We utilized 16S rRNA gene metabarcoding data from a previous study [7], which investigated how pollutant profile complexity influences the biodegradation of pharmaceuticals in wastewater microbial communities. Briefly (see details in the original study), 96 batch cultures of 20 ml synthetic wastewater were prepared following OECD guidelines and inoculated with 1% homogenized wastewater from a membrane bioreactor. Five pharmaceuticals—caffeine, atenolol, enalapril, paracetamol, and ibuprofen—were introduced in all 32 possible single and combined treatments at 100 mg/L, each in triplicate alongside abiotic controls. The cultures were incubated for 11 days at 22 °C under continuous shaking. At the end of the study, samples were destructively sampled, and community composition analyzed with 16S rRNA metabarcoding (SRA archive under ID PRJNA1041291). This study focuses on single pollutant treatments (enalapril, atenolol, caffeine, ibuprofen, paracetamol) and the controls (no pollutant).

### Pathogen Detection Pipeline

To assess which bacterial pathogens were affected by pollutant complexity, we applied the MBPD pipeline (“Multiple Bacterial Pathogen Detection”) [8]. MBPD uses 16S rRNA gene amplicon sequences as input and aligns them against a curated database of 1,986 pathogen species (72,685 full-length 16S sequences across animal, plant, and zoonotic pathogens) using UCLUST for classification. Differential abundance of abundant pathogens between each pharmaceutical compared to the control was evaluated with DESeq2 a [9]. The full code used in this study is available on github (marcel29071989).

## Results

Spiking with pharmaceuticals led to significantly higher pathogen richness and relative abundance compared to pollutant-free samples (Figure 1). The relative abundance of all pathogens exhibited a significant increase, rising from 24% ± 3 to 34% ± 1. Concurrently, the richness of all pathogens increased markedly, from 21 ± 8.1 to 122 ± 7.5 (Fig. 1a). For animal pathogens, the relative abundance increased from 20% ± 5 to 25% ± 1, while the richness rose substantially from 14.3 ± 4.9 to 82.9 ± 5.7. (Fig. 1b). Similarly, zoonotic pathogens demonstrated a clear upward trend. Their relative abundance increased from 4.3% ± 2 to 8% ± 1, and their richness expanded from 6.0 ± 2.5 to 26.4 ± 1.12 (Fig. 1c). Finally, plant pathogens also showed a notable increase. Their relative abundance rose from 1% to 6% ± 0.6, and the richness of plant pathogens increased from 0.7 ± 0.67 to 12.9 ± 0.9. (Fig. 1d).

**Fig. 1.**
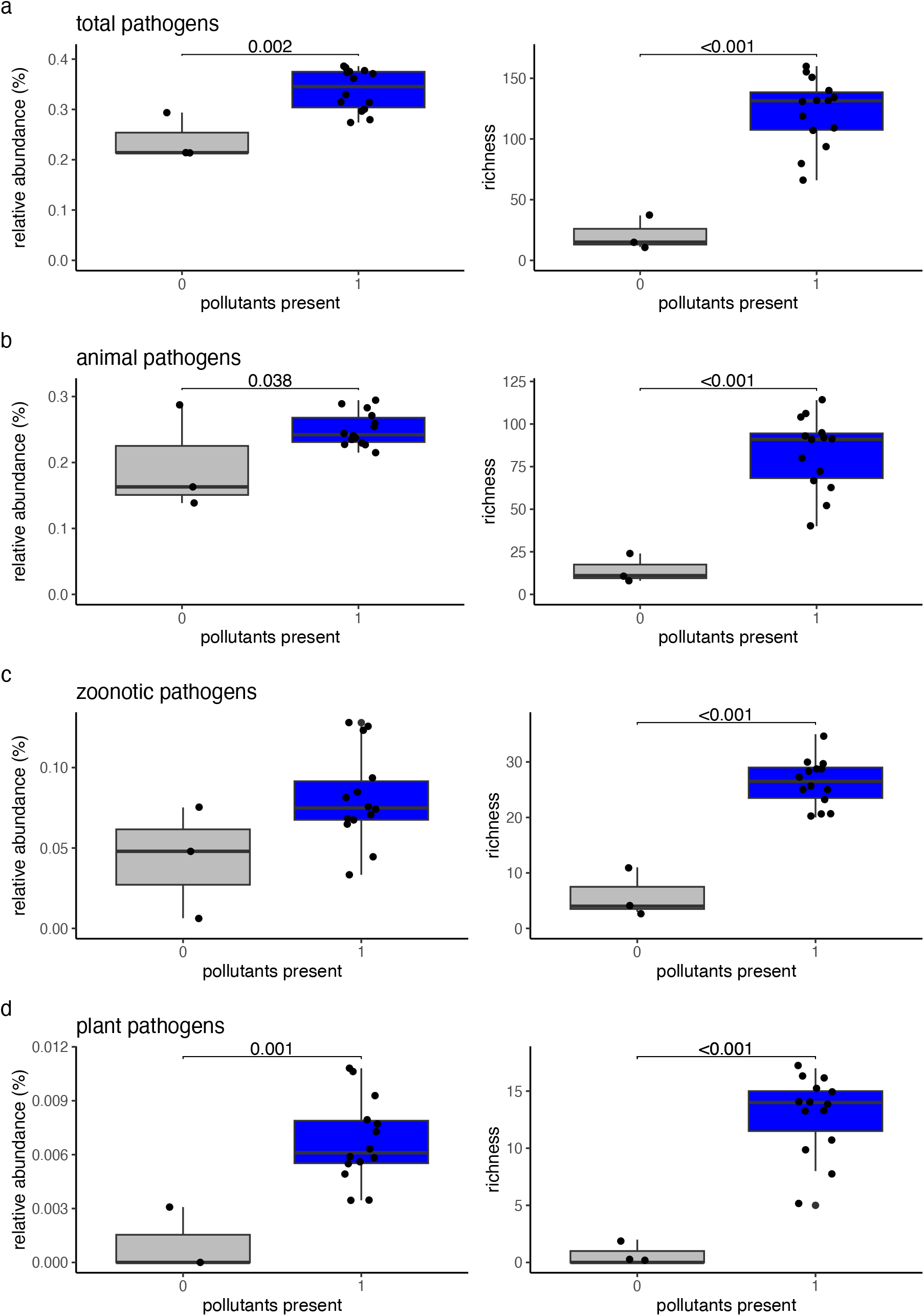
Influence of the presence (1) or absence (0) of pharmaceutical pollutants on relative abundance and richness of pathogens for (a) total pathogens, (b)animal pathogens, (c) zoonotic pathogens and (d) plant pathogens.

Furthermore, we conducted a detailed analysis to determine which specific pollutants were driving the proliferation of different pathogen groups. Our results revealed that the pooled relative abundance of all pathogens was significantly influenced by pollutant identity (Table 2), with the greatest effect observed for atenolol, increasing from 24 % to 37 % (Fig. 2a). Atenolol had an even more pronounced effect on pathogen biodiversity, with total richness rising from 21 to 144 (Fig. 2a). When separating the data per functional group, we found that the relative abundance of animal pathogens was significantly impacted by all pollutants, again with the strongest effect for atenolol, which caused an increase from 19 % to 28 %. The richness of animal pathogens showed a pronounced increase under atenolol exposure, from 14.3 to 101.0 (Fig. 2b). For zoonotic pathogens, the relative abundance was also significantly affected by all pollutants, with the highest effect observed for enalapril, rising from 4 % to 11 %. In contrast, the richness of zoonotic pathogens was most strongly influenced by paracetamol, increasing from 6 to 31 (Fig. 2c). Lastly, the relative abundance of plant pathogens was significantly impacted by all pollutants, with atenolol again exhibiting the strongest effect, increasing levels from 0.1 % to 1 %. The richness of plant pathogens also showed the greatest increase under atenolol exposure, from 0.6 to 15 (Fig. 2d).

**Table 1.**
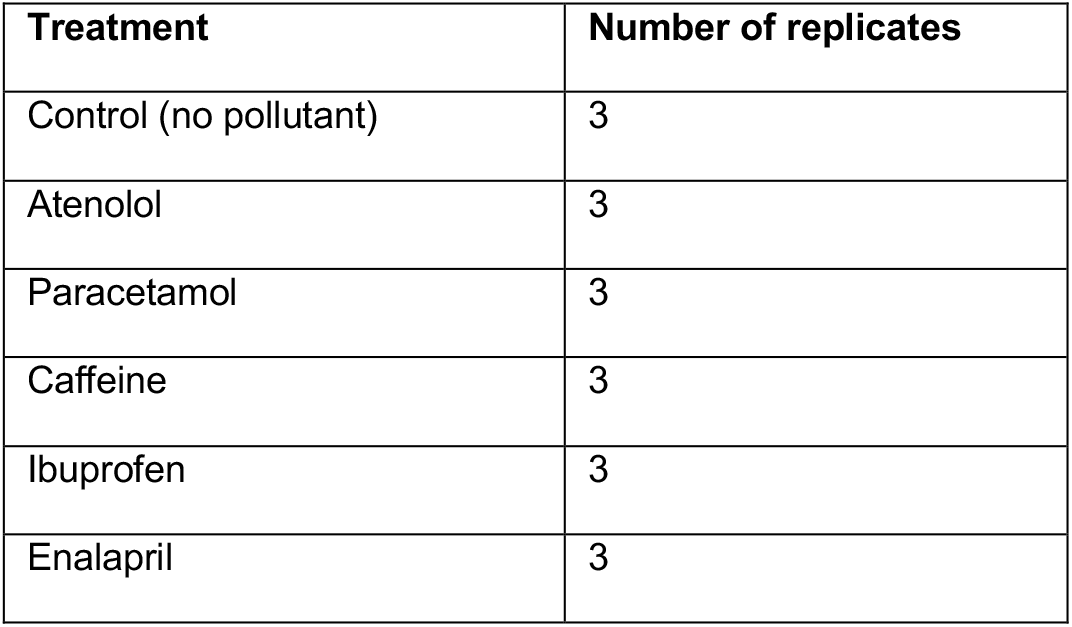
Analyzed treatments of this study.

**Table 2.** Statistical analysis of effects of presence of pollutants on pathogen abundance and richness.

| Variable | Effect of presence/absence pollutant |  | Effect of single pollutants |  |
| --- | --- | --- | --- | --- |
|  | F-value | p value | F-value | p value |
| total pathogens relative abundance | 13.52802 | 0.002237 | 3.3938 | 0.04245 |
| total pathogens richness | 35.75905 | 2.52E-05 | 10.80016 | 0.000597 |
| animal pathogens relative abundance | 5.169406 | 0.038115 | 1.644761 | 0.228165 |
| animal pathogen richness | 28.4127 | 8.41E-05 | 9.218221 | 0.001176 |
| zoonotic pathogens relative abundance | 4.110926 | 0.060758 | 2.020081 | 0.153822 |
| zoonotic pathogen richness | 57.04626 | 1.74E-06 | 18.82017 | 4.66E-05 |
| plant pathogens relative abundance | 15.92544 | 0.001182 | 10.5235 | 0.000668 |
| plant pathogen richness | 37.24687 | 2.02E-05 | 7.68215 | 0.002486 |

**Fig. 2.**
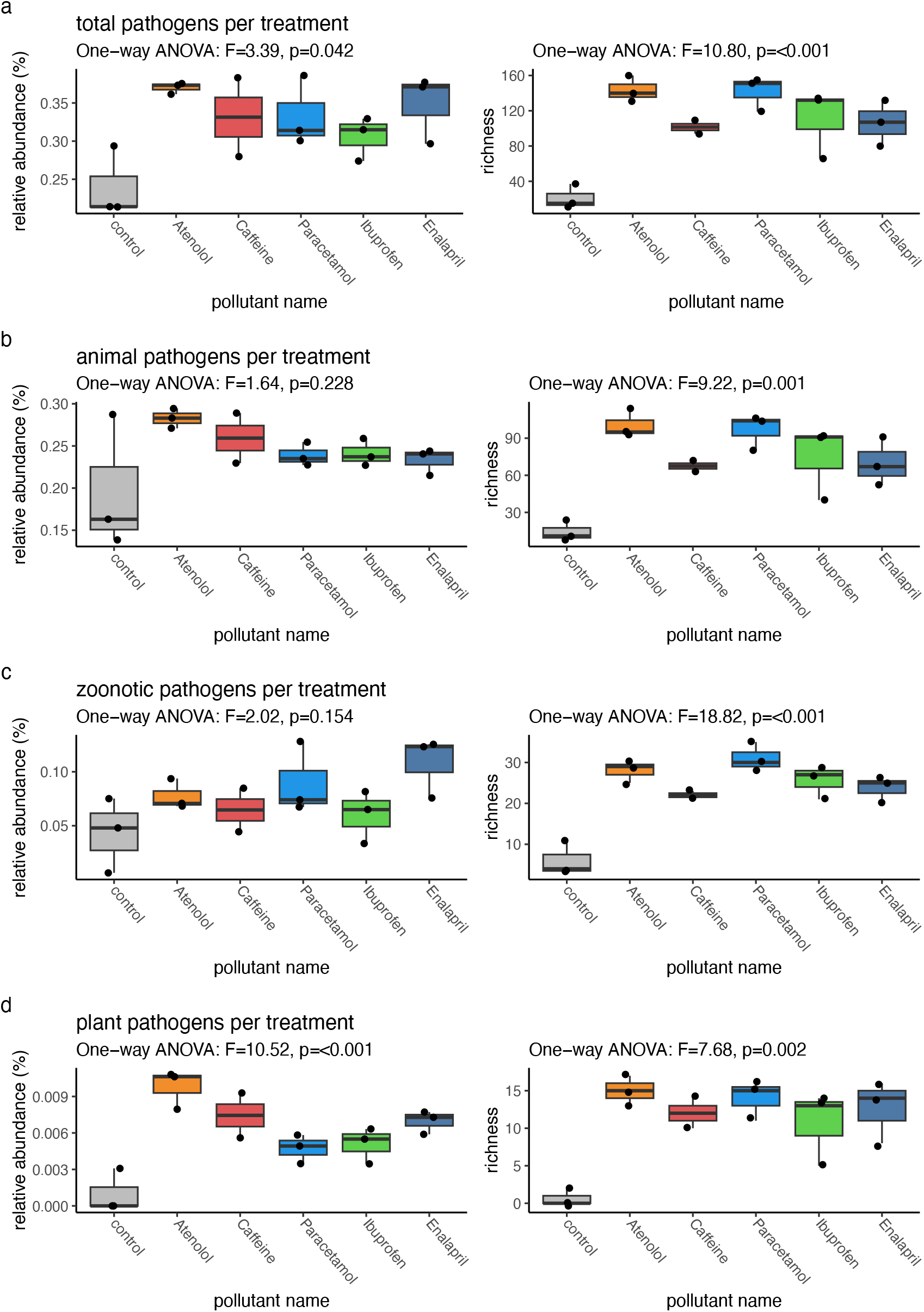
Overall effect of single pharmaceuticals on the relative abundance and richness of pathogens. **(a)** Effect of single pharmaceuticals on total pathogens abundance and richness **(b)** Effect of single pharmaceuticals on animal pathogens abundance and richness. **(c)** Effect of single pharmaceuticals on zoonotic pathogens abundance and richness. **(d)** Effect of single pharmaceuticals on plant pathogens abundance and richness. The pharmaceutical pollutants were ibuprofen, atenolol, paracetamol, enalapril and caffeine, respectively.

To identify which pathogens were significantly affected in their abundance by individual pollutants, we performed a DESeq2 differential abundance analysis. The results revealed clear pollutant-specific effects on microbial taxa. Atenolol exposure led to significant increases in *Elizabethkingia miricola, Eciguobacterium sibiricum*, and *Acinetobacter lwoffii* (Fig. 3a). Similarly, ibuprofen significantly enriched *Chryseobacterium indologenes, Flavobacterium psychrophilum*, and *Empedobacter brevis* (Fig. 3b). Paracetamol exposure predominantly affected *Acinetobacter junii, Vibrio cholerae*, and *Empedobacter brevis* (Fig. 3c). In contrast, enalapril had a more selective impact, primarily increasing the abundance of *Chryseobacterium indologenes* and *Flavobacterium psychrophilum* (Fig. 3d). Finally, caffeine showed a significant effect only on *Chryseobacterium indologenes* (Fig. 3e). Collectively, these results highlight that different pollutants selectively modulate the abundance of distinct pathogenic taxa, with atenolol and paracetamol exhibiting the broadest effects.

**Fig. 3.**
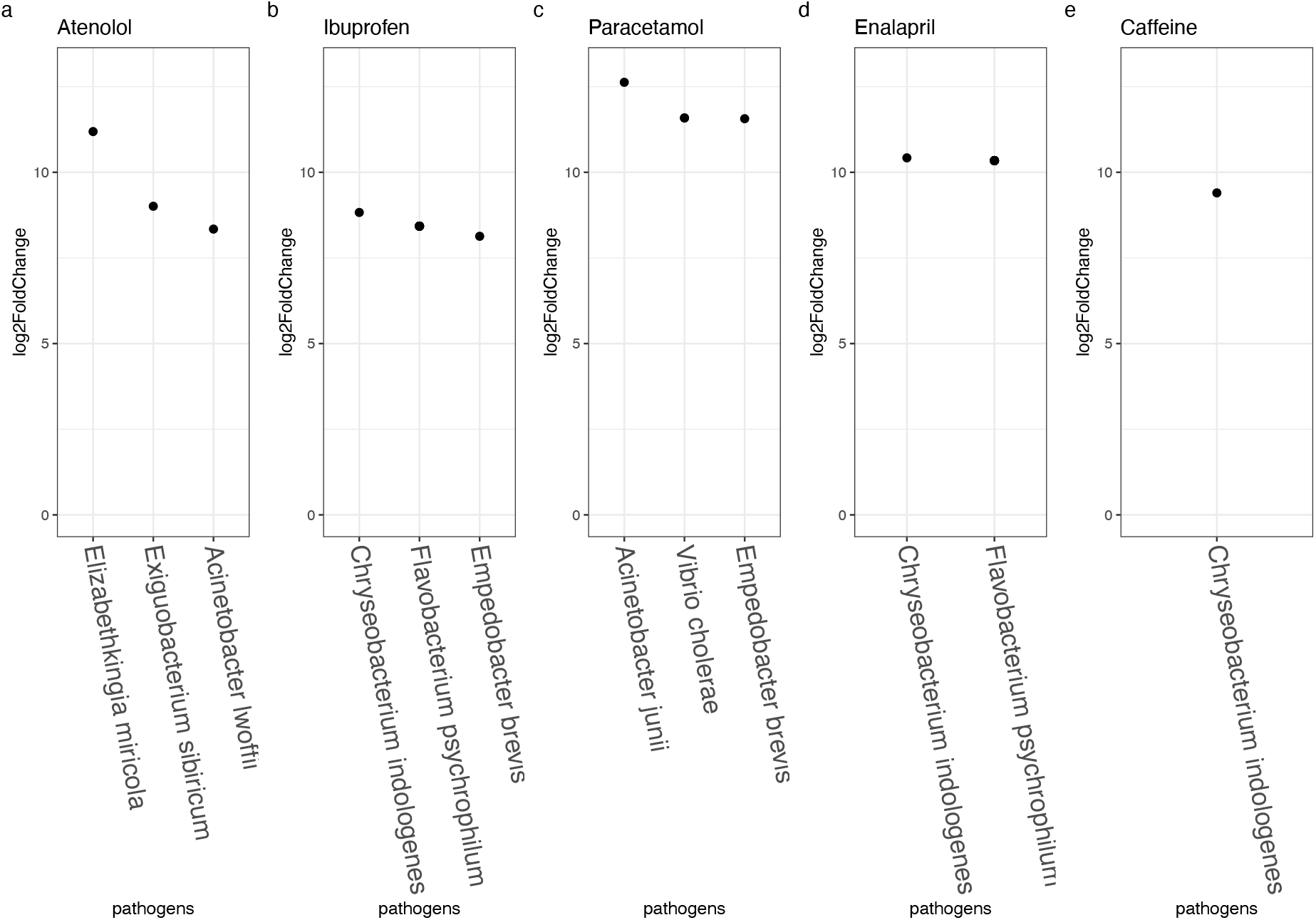
Effect of single pharmaceuticals on the presence of pathogens significant higher compared to no pharmaceutical present. **(a)** Effect of atenolol on the abundance of pathogens **(b)** Effect of ibuprofen on the abundance of pathogens **(c)** Effect of paracetamol on the abundance of pathogens **(d)** Effect of enalapril on the abundance of pathogens **(e)** Effect of caffeine on the abundance of pathogens.

## Discussion

While numerous pharmaceuticals enter wastewater treatment plants, the consequences of these compounds for the spread of pathogens remain largely unknown [8, 10–12]. Many of these pharmaceuticals, including painkillers, anti-inflammatory drugs, antidepressants, and cardiovascular medications, are not designed for antimicrobial purposes, yet they are widely consumed worldwide and frequently excreted or discarded into wastewater [5]. Their chemical persistence and resistance to degradation allow them to interact extensively with microbial communities and co-occurring pollutants, creating subtle but significant ecological pressures in wastewater systems [12]. These substances are chemically persistent and difficult to degrade, which allows them to interact extensively with microbial communities and with each other [7]. Their presence highlights an often-overlooked aspect of wastewater pollution management and public health protection. The ecological pressures they impose can suppress beneficial microbes, shift community composition, and in some cases favor the survival or proliferation of pathogens [11]. Our analysis clearly showed that these effects extend across all kinds of pathogens - zoonotic, animal, and plant - significantly increasing both their abundance and richness in wastewater environments.

One key finding of this study was the strong increase in plant pathogens under the influence of pharmaceutical residues in wastewater. Plant pathogens are often neglected in wastewater risk assessments, yet they have major implications for food security. Many agricultural systems rely on water sources that may carry wastewater-derived microbes, and chronic exposure to pharmaceutical compounds can reshape microbial communities in ways that favor phytopathogen survival and proliferation. For example, *Ralstonia solanacearum*, a highly destructive soil- and water-borne pathogen, is readily disseminated through contaminated irrigation water and is recognized as one of the top ten bacterial threats to global harvests [13]. Pharmaceutical-driven alterations in microbial dynamics can thus promote survival and spread of devastating pathogens: Several of the taxa most enriched under pharmaceutical exposure include opportunistic or high-risk pathogens. For example, *Vibrio cholerae*, promoted by addition of Paracetamol, is a major human pathogen responsible for cholera outbreaks[14]; *Acinetobacter lwoffii* and *Acinetobacter junii*, promoted by Atenolol and Paracetamol, are opportunistic pathogens increasingly associated with nosocomial infections and antibiotic resistance reservoirs [15, 16]; and *Elizabethkingia miricola* can cause severe infections in immunocompromised individuals [17]. Meanwhile, *Flavobacterium psychrophilum*, promoted by Enalapril, is a critical fish pathogen with major aquaculture implications[18], and *Chryseobacterium indologenes* and *Empedobacter brevis* are emerging opportunistic pathogens capable of surviving in water treatment environments [19, 20]. These findings highlight that pharmaceuticals do not simply shift overall microbial diversity but actively favor the proliferation of clinically and agriculturally relevant pathogens.

Among the most concerning compounds affecting pathogens was atenolol, a widely used beta-blocker that is commonly detected in wastewater treatment plants and is notoriously difficult to completely degrade. In our observations, atenolol primarily influenced the richness and abundance of pathogenic species, shaping community composition rather than directly enhancing pathogen tolerance. Its persistence in wastewater environments allows it to interact continuously with microbial populations, and over time these interactions can favor the maintenance of diverse pathogen reservoirs. Under such conditions, wastewater transitions from a neutral medium into an active conduit for pathogen persistence and dissemination.

In conclusion, the interplay between pharmaceutical pollution and pathogen ecology demonstrates that chemical and biological hazards in wastewater are deeply interconnected. Even seemingly low-risk pharmaceuticals can act as drivers of biohazard conditions, fostering the survival, proliferation, and dissemination of pathogens that threaten both public health and food security. Effective wastewater management and risk assessment must move beyond traditional chemical and microbial monitoring to evaluate pharmaceuticals for their potential to induce biological hazards, ensuring the protection of both environmental and human health.

## Data accessibility

All datasets and metadata are available on the GitHub repository Marcel 29071989 (https://github.com/Marcel29071989/), and the raw sequencing data can be found on NCBI SRA archive under ID PRJNA1041291.

## Declaration of AI use

We have not used AI-assisted technologies in preparing this article.

## Conflice of interest declaration

We declare we have no competing interests.

## Funding

The work has been funded by the European Union’s Horizon Europe framework program for research and innovation under grant ID 101060625 (project NYMPHE). This work has also received funding from the Swiss State Secretariat for Education, Research and Innovation (SERI).

## Notes

### Competing Interest Statement

The authors have declared no competing interest.

